# A Quantitative Look at Mitochondrial DNA in Genealogy

**DOI:** 10.1101/2025.02.19.638439

**Authors:** Robin W. Spencer

## Abstract

Mutations occur throughout the 16,569 basepair human mitochondrial genome and most are under strong purifying selection. Archaeologically and genealogically useful mutations, in contrast, are neutral and remain stable for centuries. In new results, multiple lines of evidence show that the latter appear on average every 2900 years and the effective genealogical chromosome is limited to about 4800 basepairs, 28% of its physical length. These mutations are already three-fold oversampled in the haplotree; all those in protein-coding genes are translationally synonymous. This evidence offers a limited prospect that additional testing can significantly increase the branching or temporal resolution of the haplotree.

## Introduction

Human mitochondrial DNA codes for enzymes and RNAs essential to life, and so the literature of its mutations is dominated by studies of their impact in disease and viability. But the disciplines of archaeology and genealogy look at mtDNA in a completely different way, being interested in neutral mutations that may be stable for millennia and thereby provide markers for maternal ancestry.

The quantitative study of mtDNA for genealogic purposes suffers in comparison to the data-rich world of Y DNA. Several questions arise from that comparison:

1. We have well-founded tMRCA (time to most recent common ancestor) values for all branches of the Y haplotree. Can we do something similar for the mitochondrial tree?
2. Thanks to extensive consumer next-generation sequencing of Y DNA, the average interval between single nucleotide polymorphisms (SNPs) is about 84 years, so many testers’ paternal DNA ancestry overlaps with their traditional paper ancestry. What is this interval for mt SNPs?
3. Most Y mutations are unique; only a few occur two or more times. But mtDNA is much smaller, so do replicate mutations appear more often? What are the implications for haplotree structure?
4. What is the effective size of the genealogic mt genome? It must be somewhere between its physical size (16,659 basepairs) and its hypervariable regions (about 700 bp). What are the implications?

The logical paths to the key findings of this work – the average years between SNPs and effective size of the genealogically useful mtDNA genome – are shown in Figure 1. These are derived from independent sources not previously exploited, namely ancient autosomal DNA and the statistics of replicate mutations. This report follows the roadmap, first with the assignment of tMRCAs to the haplotree which leads to the interval between mutations. Then functional biology and replicate mutation statistics give an effective size of the genealogically relevant genome. A date for humanity’s most recent common female ancestor and mtDNA mutation rate fall out as consequences. The overall result is a self-consistent picture with blunt implications for genealogy.

**Figure 1.**
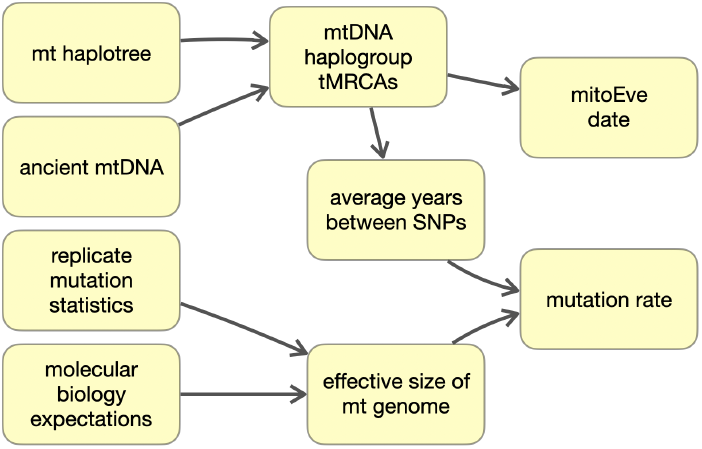
Roadmap for this research, showing how data sources and inferences are logically connected

### Updating the mt Haplotree

Assigning a tMRCA date to every haplogroup in entire mt haplotree has not been done before but is necessary for modeling and migration mapping. The task is made more complex by published mtDNA mutation rates that vary widely and differ across the chromosome^1^, as well as oversampling that results in reversions (“back mutations”) both known and invisible. Among the major sites that post haplotrees, Phylotree and Family Tree DNA (FTDNA) do not post dates and YFull dates are incomplete. Supplementary table S5 of Behar *et al* 2012^2^ is over a decade old but the most complete published list of mtDNA haplogroups and their tMRCAs.

My pragmatic approach proceeds first by mapping the Behar haplogroups to those in the current FTDNA haplotree; of the 2841 in Behar, 2692 match among the 5469 in FTDNA^3^. tMRCAs for the remaining 51% are interpolated proportionate to the number of mutations between markers, after Adamov^4^. Leaf nodes without dates are interpolated to the present day. A consistency check assures that no child node is older than its parent; the 41 cases found (0.7%) are force-fit based on the grandparent node’s date. The result is strongly correlated to the Behar dataset (Figure 2, top) and inspection of the outliers shows these to be long unbranched lineages; for example L1→L1b→L1b1 with a 120,000 year gap across two nodes, or F1c1 with 20,000 years to a leaf node.

**Figure 2.**
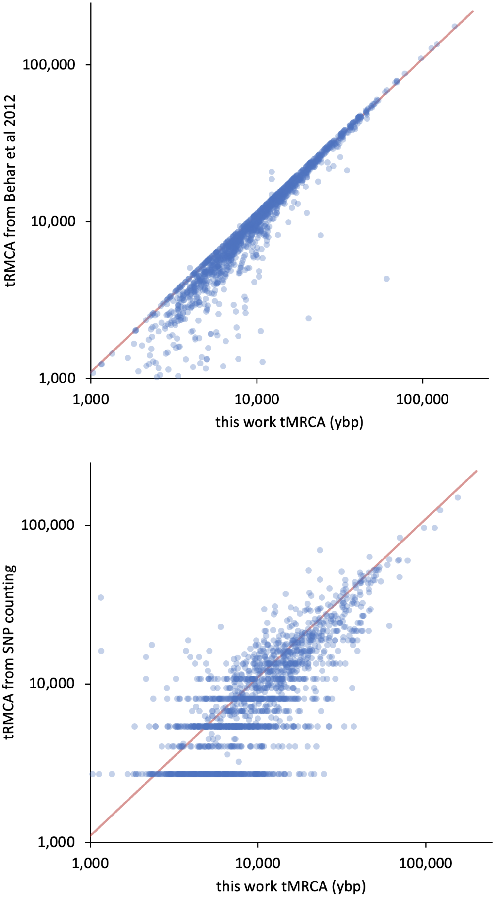
Dots are nodes on the mtDNA haplotree. Since tMRCAs here are interpolated from Behar 2012, their correlation is excellent (top) and demonstrates how little the haplotree has changed. The correlation with simple mutation counting is more scattered (bottom).

The lower chart in Figure 2 shows that simple mutation-counting, in which tMRCA is estimated as (average over all branches of mutations from the present day)×(year-per-mutation)^4^, is well correlated but with higher variance. The strong horizontal bands show the discrete effect of low mutation numbers for recent haplogroups (1 mutation from the present always gives tMRCA 2900 years, 2 give 5800 years, etc.).

Behar 2012 also gives error estimates for the 2841 haplogroups with tMRCA values. As shown in Figure 3, these largely follow the square root expectation for independent events, thus relative error decreases for ancient markers since they have more mutations to the present.

**Figure 3:**
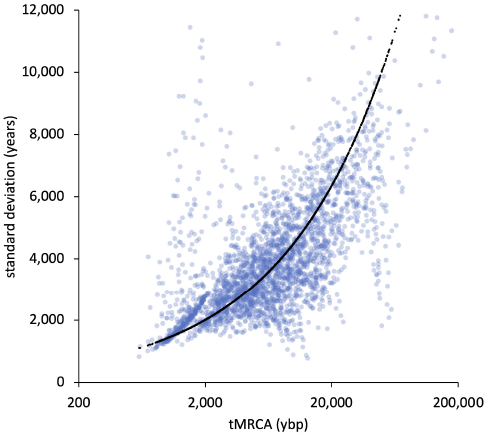
Blue: Behar haplogroups, black: 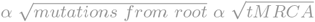

### Calibrating tMRCAs with Ancient DNA

The 2692 haplogroups with tMRCAs in Behar are a good basis for extension to the two-fold larger set of haplogroups in the current FTDNA mt haplotree. But such literature tMRCA values are ultimately based on a value for the mtDNA mutation rate. There is a large and serious literature on this with overall rates ranging from 1.3 to 4.3 × 10^−8^ mutations/site/year^5^, and these are ultimately tied to agreed historic events such as the human-chimpanzee split, major migrations between regions, or isolation of island populations. Given the 3-fold disagreement among published experts, it’s worth having a look at another very large and independent source of date information: ancient human DNA.

The concept is sketched in Figure 4, where we have three ancient samples of haplogroup “blue” and three “green”; these haplogroups are also shown by colored dots on the haplotrees above. The idea is to “stretch” the haplotree on its time axis by sliding its root (mitochondrial Eve, or haplogroup RSRS) left or right (i.e. tweaking the mutation rate) – and all markers under it, proportionately, since the time axis is linear – until the blue and green markers line up with the oldest examples of each set of ancient remains.

**Figure 4.**
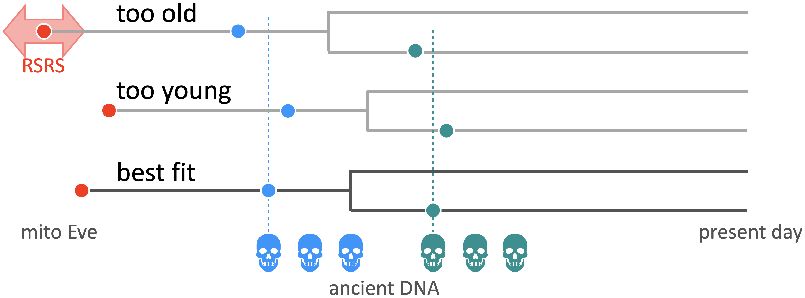
Rescaling the mt haplotree to match ancient DNA results. Time is the horizontal axis for three haplotree scenarios, in which the date for root RSRS is varied to align mutations (blue and green dots) with the oldest physical example of each (skulls).

Ancient DNA cannot be used directly or naively because the great majority of samples have assigned haplogroups that appeared thousands of years before the ancient person’s lifetime (see Figure 2 at) for example it is not unusual to have a skeleton in a medieval cemetery with a Neolithic haplogroup (Y or mt), either because a limited DNA test was done or because we have no known branches from the early haplogroup and so cannot assign a more recent label. Since we cannot say that a given ancient person was known to have lived close to the time when their DNA marker first arose, we have to estimate this threshold by looking at distributions.

Fortunately the high time-resolution of Y DNA SNPs supplies a calibrating case study. At the left in Figure 5 is the distribution of Jari Kinnunen’s FAR statistic^7^ for 5469 ancient sites with Y DNA haplogroups from the Quiles and Reich datasets. FAR cannot formally be less than one – a person cannot have a DNA marker before that marker first appeared – but with the intrinsic statistical uncertainty of haplogroup dates, there will always be some samples with FAR *<* 1. Because Y SNP dates and ancient sample radiocarbon dates are accurate and independent, this dataset sets a precedent for the left tail of the distribution which is that 3.9% of the samples have FAR *<* 1 (in dark blue).

**Figure 5.**
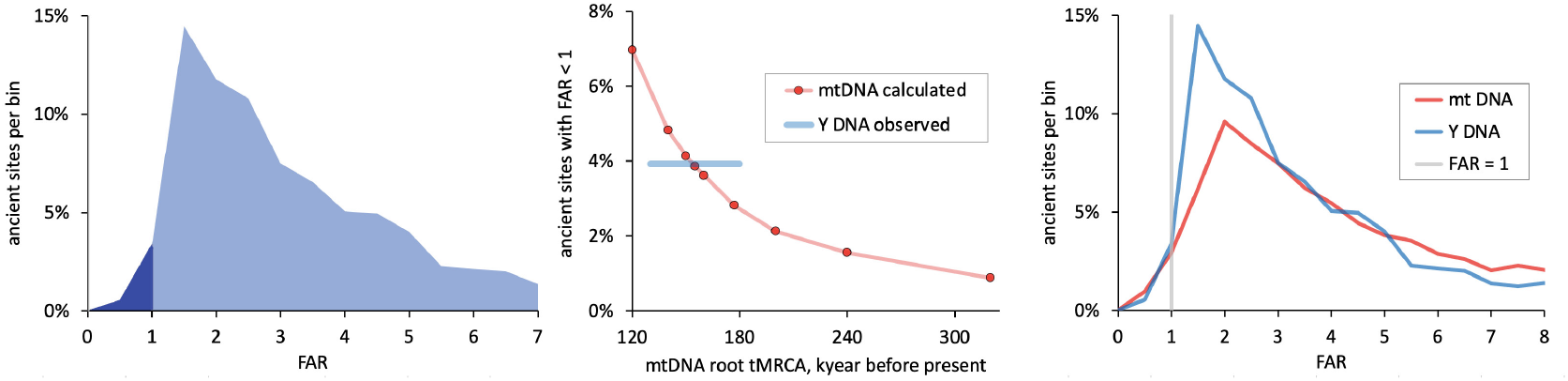
Distributions of Kinnunen’s FAR statistic. Left: FAR for 5469 ancient DNA samples with Y haplogroups; impossible values (FAR ¡ 1) in dark blue. Center: Percent of samples with mtDNA FAR ¡ 1 as a function of mtDNA root tMRCA, showing where mtDNA intersects the Y DNA observation. Right: Overlap of the Y and best-fit mtDNA distributions.

Each red dot in the center chart of Figure 5 is a full recalculation of the mt haplotree, “stretched” as the root tMRCA varies from 120,000 to 320,000 ybp. This slides the mtDNA FAR distribution of 16,698 ancient samples back and forth, and it matches the Y DNA value (3.9% with FAR *<* 1) at tMRCA_RSRS_ = 155,000 ybp. This mt FAR distribution is overlayed with that for Y DNA in Figure 5, right. This root date is a satisfying result in the middle the literature values shown in Figure 6.

**Figure 6.**
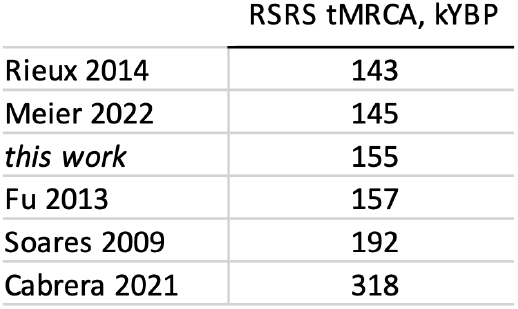
tMRCA values for root RSRS from the literature and this work.

Behar 2012 is the largest literature dataset of mt haplogroup tMRCAs, though there are several others, shown in Figure 7. As expected from the agreement with tMRCA_RSRS_, the FAR-calibrated results are in good agreement with all but Cabrera 2020, which has tMRCAs about twofold greater than any of the other datasets.

**Figure 7.**
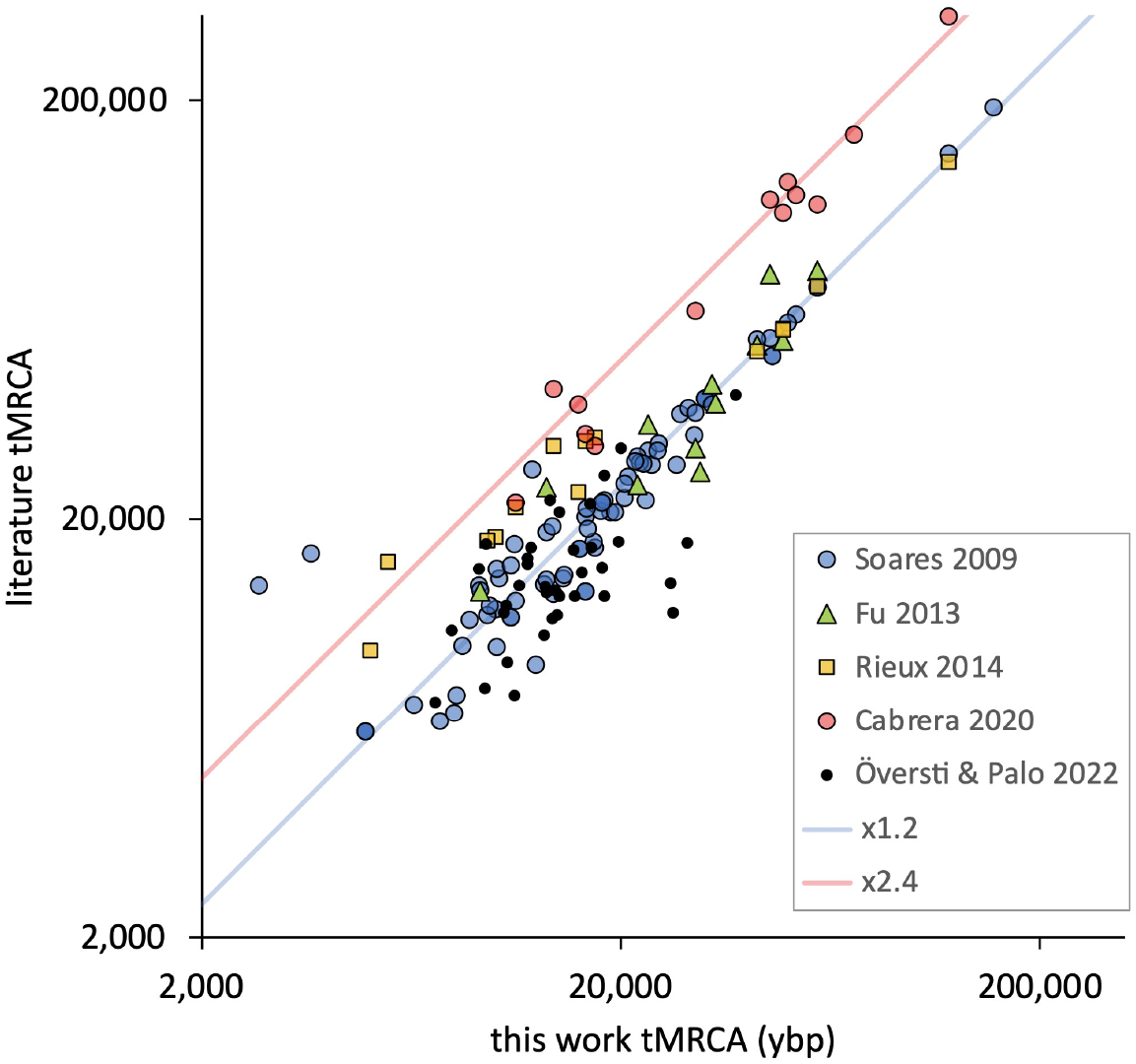
Comparison of literature tMRCAs with this work; each dot is a haplogroup. Same as Figure 2, top, but for other sources. No one source covers the range of the entire haplotree, and none is as complete as this work with 5469 haplogroups.

**Figure 8.**
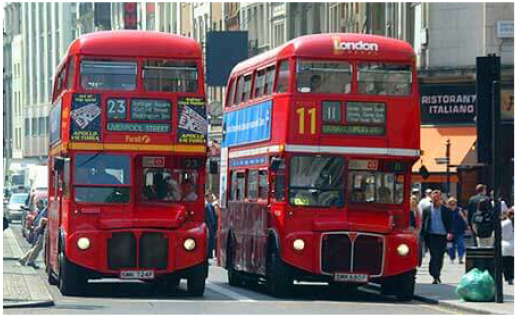
The metaphor for exponentially distributed inter-event times.

My values for tMRCAs and estimated SNP formation date are available for the entire mt haplotree at [SNP Tracker](https://scaledinnovation.com/gg/snpTracker.html): select mtDNA (♀), enter a haplogroup, and select the SNPs tab.

### Years Between Mutations

With calibrated dates assigned to the mt haplotree, the second item on the roadmap is the estimation of the years between mutations^8^ for mtDNA. This is an intuitive way to express the time resolution of a haplotree – for Y DNA the current value for BigY testers is about 84 years, so we know to expect a new SNP on a lineage about every three generations.

There are three ways to estimate the average years between mutations: an easy way, an uncertain way, and a statistical way. The easy way is to count all of the mutations between present-day testers and the root of the haplotree (Mitochondrial Eve or RSRS) and just divide that into the years since Mito Eve lived. We can count mutations in two ways^4^, getting values of 49-56 mutations between Mito Eve and today (Behar 2012 finds 57.1); that gives years-per-mutation = (155,000 years)/(49 to 56) = 1950 to 3140; we’ll use a value of 2900 which is consistent with the effective genome length (below).

The second way works well for Y DNA but is uncertain for mtDNA: The average interval between mutations is then easily calculated as 1/(mutation rate × basepairs tested) which, for 15 million (what BigY 700 measures) and a consensus literature mutation rate of 8 × 10^−10^ mutations per basepair per year, gives 83 years per SNP^9^.

But this calculation would be uncertain for mtDNA because we have neither an agreed-upon relevant length of the genome nor single applicable mutation rate – in fact those are values that this report hopes to put on a better footing.

The third, statistical, method has to do with the old adage about buses, “You wait for ages and then three show up all at once.” That may be annoying but it isn’t perverse – it has to be true if the buses are random and independent. Between-arrival times for such processes have an exponential distribution, which means that a few intervals will be long (“wait for ages”) and many will be short (“all at once”). The intervals will have an average (mean) time, say 10 minutes, but that’s not the peak of a bell-shaped curve but instead a point on a steep slope.

Mutations, being random and independent events, should obey the same statistics, as Y DNA demonstrates. Figure 9 shows the distribution of the years between Y SNPs in excellent agreement with the exponential expected for a Poisson process^10^. The slope is 1/(mean years-between-mutations) which for the dashed line is 1/(95 years) – a reasonable value since the Y haplotree is a mix of BigY-500 and BigY-700 datasets, which sample 10 and 15 million basepairs respectively, which give mean intervals of 100 and 84 years by the easy calculation above.

**Figure 9.**
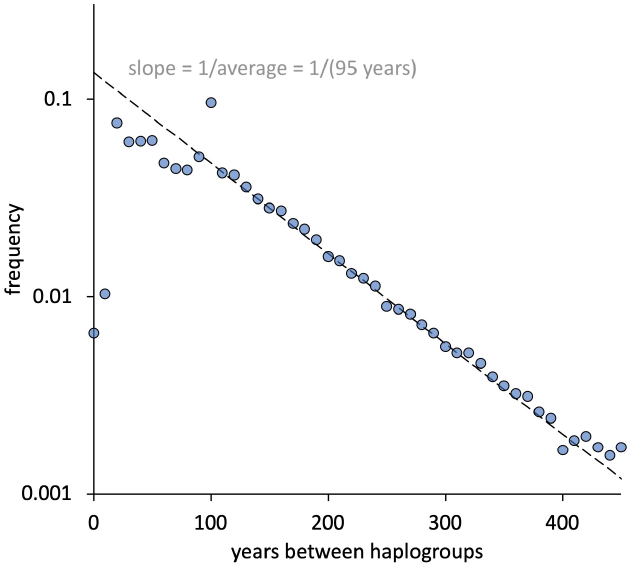
The distribution of between-mutation intervals for the 23,227 unit segments of the Y haplotree, i.e. those without intervening unlocated mutations.

Mitochondrial DNA also shows an exponential distribution in the tail of inter-mutation intervals (Figure 10) with a slope of 1/(2700 years), in good agreement with the counting method. The fall-off at low intervals probably reflects the linear interpolation used to assign haplogroup tMRCAs for leaf nodes, a form of averaging that would wash out the pattern.

**Figure 10.**
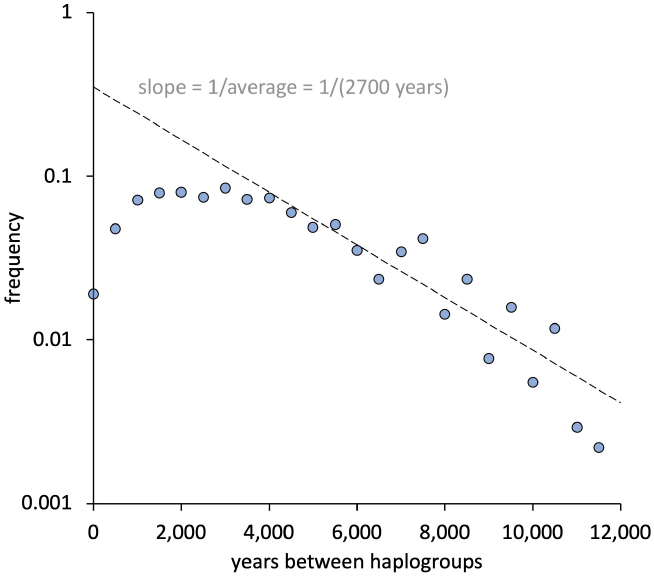
Same for the 2737 unit segments of the mitochondrial haplotree.

The point of exploring this statistical method is not to find the values for years-per-mutation which we know by simple SNP counting – the point is to illustrate with the real data that these intervals are exponentially distributed. Just like waiting for a bus, you may see a very long gap between mutations or several short gaps. Any confusion can be blamed on the introductory teaching of probability and statistics which emphasizes normal (bell-shaped, Gaussian) distributions. Many things in the real world have long-tailed distributions and we have to let go of our tacit assumption of bell-shaped behavior. Even using the term “normal” for the Gaussian distribution is problematic.

### Mapping Mutations to the Functions of mtDNA

Next on the roadmap is an estimate of the effective size of the mitochondrial genome, but first a brief review of mtDNA composition and function provides context and supportive arguments.

The human mitochondrial genome is a doublestranded circle of 16,569 basepairs of DNA that encodes 13 proteins, 22 tRNAs, and 2 rRNAs, all of which are essential for life. As shown in Figure 11, top, protein and RNA coding make of 92% of the genome. Mutations are simply named, for example G5147A tells us that a G (guanosine) at position 5417 has changed to an A (adenosine). We can look up position 5147 to see that it’s the third base in a triplet coding for threonine in the gene ND2, NADH dehydrogenase subunit 2. Mapping all mutations in the mtDNA haplotree (or Phylotree, which is very similar) by function gives Figure 11, bottom, where HVS = hypervariable sequence and CR = control region (which includes the three HVS’s). Multiplying through shows that about half of all observed mutations occur in protein coding genes, a third in HVS, and most of the rest in RNA coding genes.

**Figure 11.**
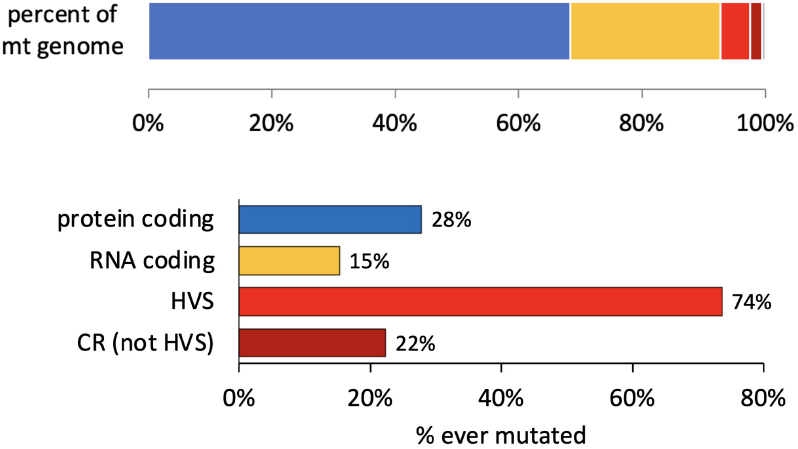
Top: size of mtDNA regions. Bottom: Genealogically relevant mutations in mtDNA by region.

Figure 12 shows the positions of all mutations in the haplotree; the bushy hypervariable segments are clearly visible, the RNA genes are relatively bare, and otherwise mutations appear evenly distributed across the protein-coding genes. Mapping the latter against the genetic code reveals something very interesting: *all* of the observed mutations in the protein genes are *synonymous*, which is to say that these mutations will not change the amino acid in the translated protein^11^.

**Figure 12.**
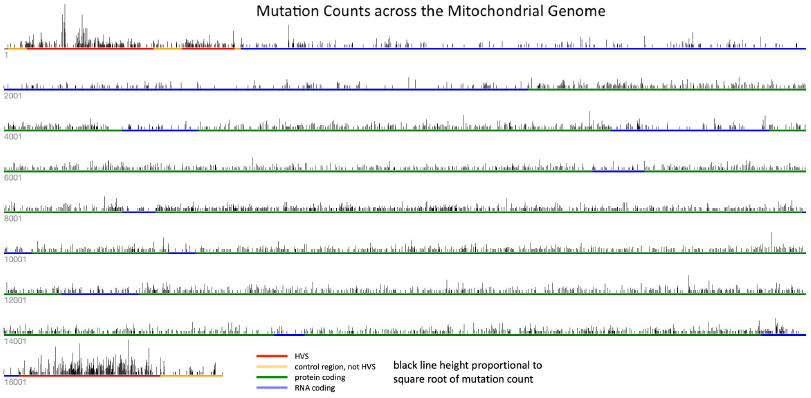
Visualization of genealogically relevant mutations in mtDNA, color coded by function. Each vertical black bar is one location with height proportional to the square root of the number of mutations observed there. Click to enlarge.

**Figure 13.**
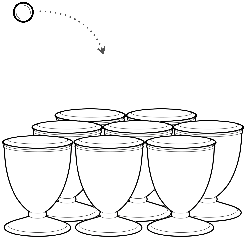
A statistical metaphor for random, independent mutations (balls) at locations in DNA (urns).

Counting up all possible mutations from one triplet to another shows that 132 of the 444 possibilities, 24%, are synonymous. Figure 11 bottom shows that 28% of the protein-coding positions have ever been observed to mutate (all of which are synonymous). This cannot be coincidental and the inference is significant in predicting the future of mtDNA testing:

All possible synonymous (silent) mutations in the protein genes are already observed in the haplotree, and no mutation occurs in a non-synonymous position, which would be under purifying selection and genealogically invisible. Since protein genes account for 68% of the mt genome, its effective length can at most be half of its physical length.

It must be smaller yet since mutations in only 15% of the RNA-coding genes are seen in the haplotree, suggesting that a larger fraction of RNA positions are under selection, which would be expected for all binding and secondary structural elements. Perhaps only solvent-facing external loops in RNA would be truly silent – it would be interesting to map the observed positions against known RNA structures. This reasoning allows a first approach to the number of stable and observed mutations in genealogically-relevant mtDNA: from the haplotree we have 13,236 mutations of which 4626 are unique, a 2.9-fold oversampling (these 4626 occupy 4128 positions in the chromosome: the control and HVS region plus 28% of protein genes plus 15% of RNA genes). If the oversampling is complete, the effective mtDNA genome allows only 4626 mutations. We wouldn’t find other mutations because they’d be under selection pressure, making those mitochondria (or cells, or embryos) inviable and invisible, certainly over millennia^12^.

### Tossing Balls into Urns and Counting Repeated Mutations

This is an important conclusion so it’s worth approaching from another direction with different data. If mutations are independent, then their statistics should be the same as statisticians’ favorite pastime: tossing balls blindly and randomly into urns. Thought experiments set the groundwork:

- If we toss 10 balls into 10,000 urns, we’re very unlikely to have more than one ball per urn. We’ll probably end up with 10 urns having one ball each and 9,990 empty urns. This system is very *unsaturated*.
- If we toss 1000 balls into 10 urns, none will be empty and the rest will have an average of 100 balls each, in some nice bell-shaped distribution. This system is oversampled and *saturated*.
- If we toss four balls into four urns, we probably won’t get one in each, nor all four in one urn. More likely is something in between like one urn with two balls, two urns with one each, and one urn left empty.

Such ball-tossing gives rise to Poisson distributions^13^ where the mathematics is well understood. This means that we can treat ball-tossing as an experiment to find the total number of urns when the empties are invisible to us and we can see only those urns with balls in them. The parallel to mutations is direct: if we find that all mutations occur in different locations (one ball per urn), then the genome must be very large, or if all mutations pile up about equally in a smaller number of locations, those locations are the whole genome, and if there’s a distribution in between, we can use Poisson math to estimate the total genome size.

Turning first to Y DNA, at this writing the Y haplotree lists 659,485 mutations, of which 616,709 (93%) are seen only once, 17,888 (5.4%) are seen twice, and 2010 three times, as shown in Figure 14. The blue curve shows what a Poisson result tells us to expect when tossing 660,000 balls into 11,000,000 urns; it’s not a bad fit for the first two points and 11 million is a decent extrapolation to the known testing lengths of BigY (10 and 15 million). Curiously the data fit an exponential far better, though there is no basis for that in balls-and-urns statistics; there may in fact be a few subtle hotter-spots for mutation in the sampled Y genome.

**Figure 14.**
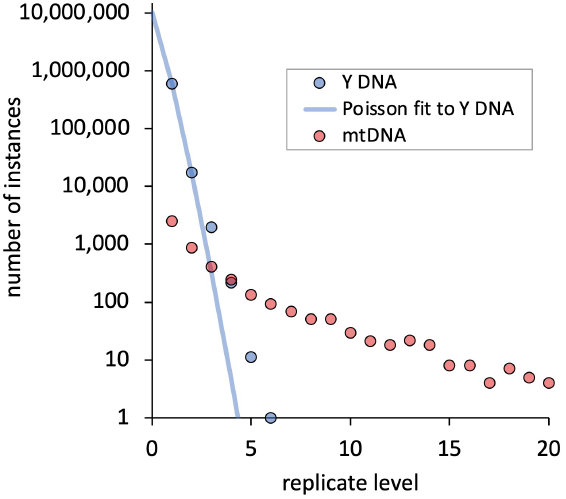
Distributions of replicate mutations. Replicates in Y DNA are rare and approximate the expected Poisson distribution. Those in mtDNA are frequent and their distribution is long-tailed.

Since the mitochondrial physical genome is a thousand-fold smaller than the sampled Y chromosome, and mtDNA is just as old as Y DNA, we expect, and observe, many more repeat mutations; Figure 14 shows that of the 4626 distinct mutations in mtDNA, 2518 (19%) are seen just once, 861 (13%) are seen twice, and so forth for a very long tail: position position 152 (specifically mutation T152C!) occurs 194 different times in the mt haplotree.

The distribution of mt replications does *not* fit a Poisson distribution. Figure 15 repeats the mt data and shows Poisson distributions for lengths of 16,569, 4626, and 702 basepairs – respectively the lengths of the whole genome, observed mutations, and hypervariable regions. None is anywhere close to the observed distribution, nor are any of their fractional sums, especially because the observed long tail goes out to 194. This is a surprise, not noted in the extensive mtDNA literature, and so worth a digression.

**Figure 15.**
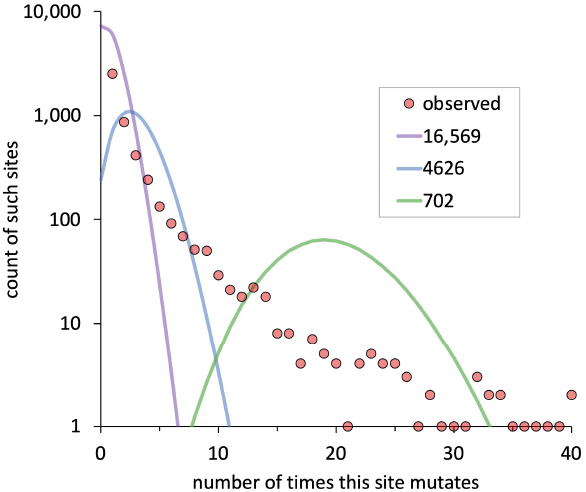
Red dots: the distribution of mtDNA replicate mutations, overlayed with (solid curves) the expected Poisson distributions for genomes of given length.

### Power Law Distributions of mtDNA Mutation Frequencies

Figure 16 shows the mtDNA mutation frequency data of Figures 14 and 15 cast as rank-frequency, and the long tailed nature of all datasets – control region, coding genes, and entire genome – is very apparent. These are distinctly not Poisson distributions, which are short-tailed (and converge to normal distributions at high N). They strongly appear to be Pareto (power law) distributions that exhibit saturation (a drop-off at the right), the result expected of a positive-feedback generative process operating in a finite domain. This is confirmed by simulations which have only one fitted parameter, alpha, the slope of the power law, with values of 2.6, 3.5, and 2.75 for control, coding, and whole regions respectively^14^; the size of each region and number of mutations are set by haplotree values. Selecting “random independent” in the simulator will show how different the observed data are from a Poisson process.

**Figure 16.**
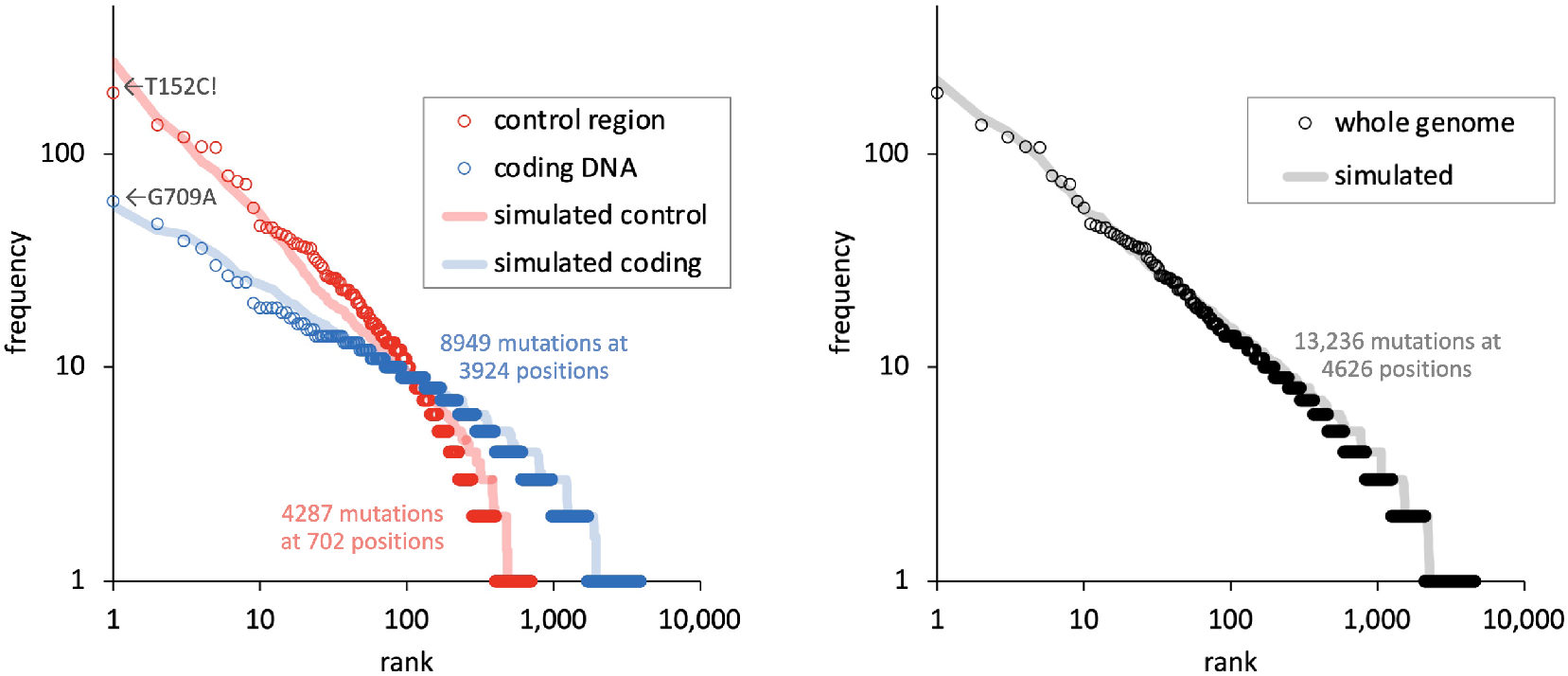
Rank-frequency distributions of mtDNA haplotree mutations by region. The most-replicated mutations have rank = 1. Solid curves are simulations for a power-law generative process in a finite domain^15,*Bagrow*^, with parameters in the text and [simulator](https://scaledinnovation.com/gg/mtDNAbagrowModel.html).

These distributions raise some interesting questions:

- Power laws are usually diagnostic of a positive-feedback generative “rich get richer” process. That makes sense for wealth, network topology, and forest fires^15^, but it’s difficult to imagine a positive feedback process (which implies memory or adjacency) for DNA mutations alone. Options and implications are discussed below.
- The means of these datasets are 2.3 mutations per location for coding DNA, 6.1 for control DNA, and 2.9 for the whole mt genome, which suggests that the control region really is 2.7-fold “hotter” than coding DNA, even when all of the non-synonymous positions are excluded. Does this reflect a biological difference, perhaps in extent or duration of protective protein coverage or exposure during replication?
- The fact that coding DNA – not just the control region – exhibits a saturating power law^15,*Bagrow*^ argues against Poisson randomness in all cases and deflates arguments specific to the control region like hot-spots or poly-cytosine domains.

As a recent review notes^16^, much early mtDNA lore has been discredited (for example anything special about oxidative damage) and there is much still to be understood. The balls-in-urns model works reasonably well for Y DNA and counting replicate mutations gives a good estimate of its effective length. Though mtDNA mutation replication statistics are clearly not Poisson, saturation is still apparent. If the process were random we can still estimate how much bigger the effective mt genome might be: for a balls-in-urns process, the probability that an urn gets zero balls is P_0_ = e^*−λ*^, where *λ* is the mean which for us is 13384/4560, so that P_0_ ≈ 0.05. Thus our observed nearly-saturated set of 4626 mutations implies a maximum of about 4626/(1 - 0.05) = 4900 possibilities for the genealogically effective mtDNA mutations. We can use 4800 as a working estimate.

### Options for Power Law Generation - a Open Problem

The appearance of mutations should be an intrinsically random Poisson process, but the observed frequencies have long-tailed distributions; therefore we must look to the larger context of mtDNA for an explanation. Options include

#### Distribution of Mutation Rates

The distributions of Figure 16, left, show that the control and coding regions have 3-fold different apparent mutation rates, yet when aggregated (Figure 16, right), the combined dataset displays a single distribution. Rather than aggregating, extrapolating backwards to fragment the genome into smaller and smaller sets of mutatable locations suggests the possibility of significantly different intrinsic mutation rates perhaps down to the basepair level. Figure 15 might then be accounted for as the sum of many – not just two or three – Poisson distributions, such that what we see reflects the distribution of mutation rates. Try the simulator with “variable mutation rates” to see that exponentially distributed mutation rates approximate a power law at early times. Other distributions could be hand-picked to get a closer result, which makes this a hand-waving, circular model.

#### Association by Survival

Figure 17 illustrates the concept. A cohort of mtDNAs exist for sufficient time for each to acquire a small number of mutations, with orange dots signifying a mutation in a non-synonymous coding position and green dots neutral mutations. By chance, some mitochondrial will have DNA with only green (neutral) mutations, even though on average orange will outnumber green by about 2.5-to-1 (the ratio of their lengths). Next a culling process occurs at the level of mitochondrial function (with the simplifying assumption of one mtDNA genotype per mitochondrion) such that any genome with an orange-dot mutation is removed from the pool. Those all-green will survive and expand to create the next generation. Some in this cohort will have no mutations, some one green mutation, and some more than one. Those latter mutations will appear to be causally connected, as if one green biases toward additional greens. But there is no actual causality between the green mutations, only an editing process that removes everything else. However whether such imposed connections between observed mutations suffice to generate a long-tailed distribution is unclear – this may be only part of a necessarily more complete model.

**Figure 17.**
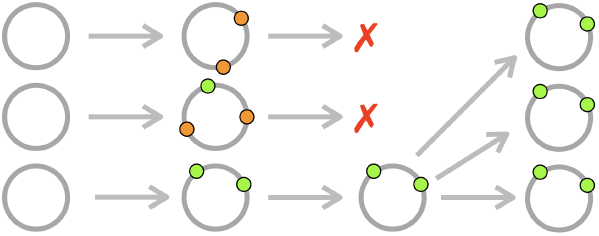
Association by Survival model. Gray circles are the mtDNA chromosome, orange dots are lethal mutations, green dots are neutral mutations.

#### Monkey Typing

The very simple (and therefore powerful) model of monkey-typing by Miller challenged more elaborate theories to explain Zipf’s observation of power laws in word lengths^15^. For Miller’s typewriter letters, substitute neutral mutations, and for the space bar, substitute harmful mutations – which could give us mtDNAs with a power law in the number of neutral mutations that they carry. This isn’t exactly what we measure in Figure 16 (the frequencies of mutation that repeat at specific locations), but it’s a generative mechanism to consider.

#### Intermittent Avalanches

The model of Newman *et al* ^15^ is also intriguing: an ensemble of independent agents (sandpiles, species) is subject to continuous perturbation over time (addition of sand grains, environmental stress) which, above a threshold, makes them subject to catastrophic removal (avalanches, extinction), and even though independent, the agents display a power-law distribution of their lifetimes. By analogy, mitochondria are independent and subject to continuous perturbation (mutations), some of which are detrimental and above a threshold make them subject to removal (mitophagy). Neutral mutations don’t trigger mitophagy and so are essentially markers for the passage of time, and may be expected to show power law distributions in their frequencies.

### mtDNA Mutation Rates

The literature values for the mutation rate of human autosomal (nuclear) DNA is in the range 0.5 − 1 × 10^−9^ *bp*^−1^*y*^−1^, and for his detailed work on Y SNP dating, Iain McDonald^9^ uses a value of 0.82 × 10^−9^ *bp*^−1^*y*^−1^. Literature values for the mtDNA mutation rate vary much more widely^5^, though most are near 2 × 10^−8^ *bp*^−1^*y*^−1^, a value that is 20-fold higher than autosomal DNA. Rieux 2014 cite a rate for hypervariable regions that’s 15-fold higher yet, though from the distributions of repeated mutations I find only a 3-fold difference (Figure 16, left).

I have deferred the discussion of mutation rates because of these unresolved differences, and because for genetic genealogy the mutation rate is a hidden parameter: what matters more are things that can be directly measured and interpreted, and from which the mutation rate can be calculated instead of the other way around. The factors are simply related:

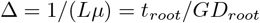

where Δ = interval between mutations (years), *L* = genome length (basepairs), *µ* = mutation rate (bp^−1^y^−1^), *t*_*root*_ = time from the present to the root of the haplotree (years), and *GD*_*root*_ = genetic distance to the root (number of mutations).

The discussion above provides values and independent justification for all but *µ*: Δ = 2900 y, *L* = 4800 bp, *t*_*root*_ = 155,000 years, and *GD*_*root*_ = 54 mutations. Solving for *µ* gives 7.7 × 10^−8^, a value that is 3-4-fold higher than those published.

This discrepancy flags a persistent problem in the literature: everyone carefully counts mutations to get a value of mutations *per genome* and then casually amortizes that over the whole 16,569 length to quote a number in mutations *per basepair*. The problem is that this simple division implies that observed mutations are equally probable across the genome, and yet we know that about 70% of the genome is out-of-bounds with bases under strong selection. I suggest that 8 × 10^−8^ *bp*^−1^*y*^−1^ is a reasonable mutation rate across the genome, but then selection (via aggressive mitophagy, both continuous and in bottlenecks, and also over genealogic timescales) edits out the 70% of wounded mtDNAs, effectively making their mutations invisible in the haplotree.

Why does this matter? Because if we persist in thinking that the entire 16,569 basepair genome is genealogically accessible, we won’t understand why the haplotree has been stalled at its modest size.

### Looking Farther Back

Some get concerned with where to “start the clock”. The RSRS sequence of Behar 2012 is a good surrogate, though of course discoveries of very early new branches like that of Maier 2022^1^ may push it back. Some authors (and bloggers) seem to assume that the root of our mtDNA (and Y DNA) haplotrees should equate to the earliest *Homo sapiens sapiens*, but the extent to which that’s true is coincidental.

Evolution is continuous and our tidy definitions of species apply only in distant hindsight. Just for fun, Figure 18 extends the correlation of genetic distance (mutation counts) vs. tMRCA to include our common ancestors with Neanderthals, Denisovans, and chimpanzees^17^. The observed difference counts are divided by two since we share common ancestry (not descent) with these other primates.

**Figure 18.**
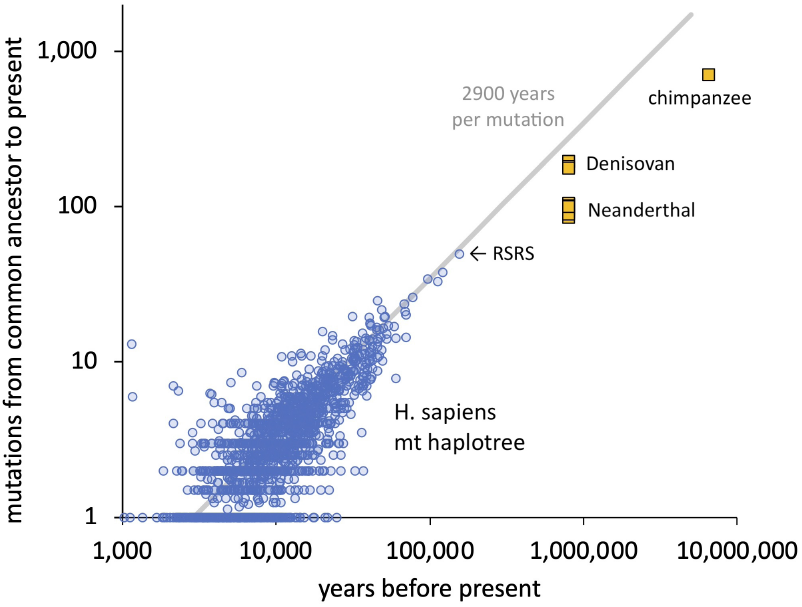
Haplotree mutations from common ancestor to present vs date of that ancestor. This extends Figure 2, bottom, to include *Homo* subspecies and chimpanzees.

The extrapolation of the linear 2900 years per mutation discussed here is not bad, though there may be some impact of reversions since the others are below the linear extrapolation. The same is seen for Y STRs. Reversions are indicated in mutation names with “!”, such as T152C!, and account for only 2.4% of all mutated positions or 2.6% of total mutations. Since they are formally invisible these mutations must be inferred by the parsimony rules of haplotree assembly and their numbers may be greater, but so far do not appear to be a major source of non-linearity – though Figure 18 hints at when correction may be necessary.

### Private Variants to the Rescue?

All of the conclusions above are based on data accessible without login or password, which is my general practice. However some early responses to this report brought my attention to private variants, namely mutations that can only be seen (at the FTDNA website) by the DNA tester. As an administrator at a large site (EnglandGB-EIJ), I have access to mutation lists for 2569 full-length mtDNA tests, and mapping these mutations against those in the public haplotree reveals the opportunity (Figure 19).

**Figure 19.**
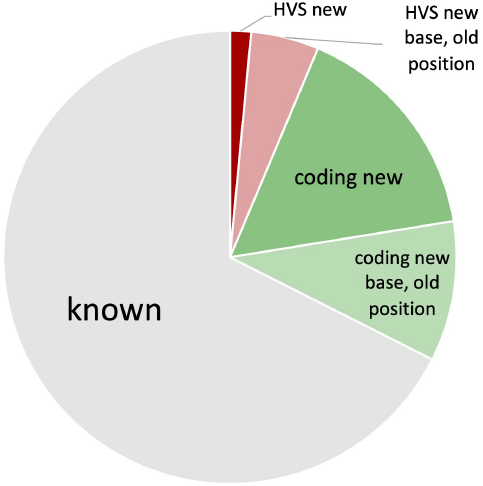
Properties of mutations seen in a sample of 2569 full-length mtDNA testers. Two-thirds are known in the current haplotree; most of the rest are in coding DNA.

Two-thirds of the mutations in this dataset are already known in the haplotree. Half of the new mutations are entirely new in the coding region, and most of the rest are new base-changes (like T→A instead of T→C) at a position already known to be mutatable^18^.

If this held up at scale, then a rough extrapolation suggests that the total known mutations could rise from 4626 to about 6100, and the average years between mutations down from 2900 to about 2200. These would be very welcome. Recomputing the haplotree including all private variants may be the most direct way to increase the resolution of mtDNA.

### Looking Ahead: the Prospects for mtDNA Genealogy

From 2018 to 2025 the size of the Y DNA haplotree grew nine-fold, a rate of 10,000 branches per year. In contrast the mtDNA haplotree has not been significantly updated since 2016: nine years at this writing. Figure 20 shows in blue the numbers of nodes in the mt haplotree from its first appearance in 1987^1^ to the present. The tree grew rapidly from 2008 to 2012, but by Phylotree build 17 in 2016 had essentially reached its current size of 5469 nodes. At about that time FTDNA offered full-length sequencing to the public and the number of mtDNA testers exploded (Figure 20, red), growing ten-fold from 2014 to 2024 but notably without an effect on tree size.

**Figure 20.**
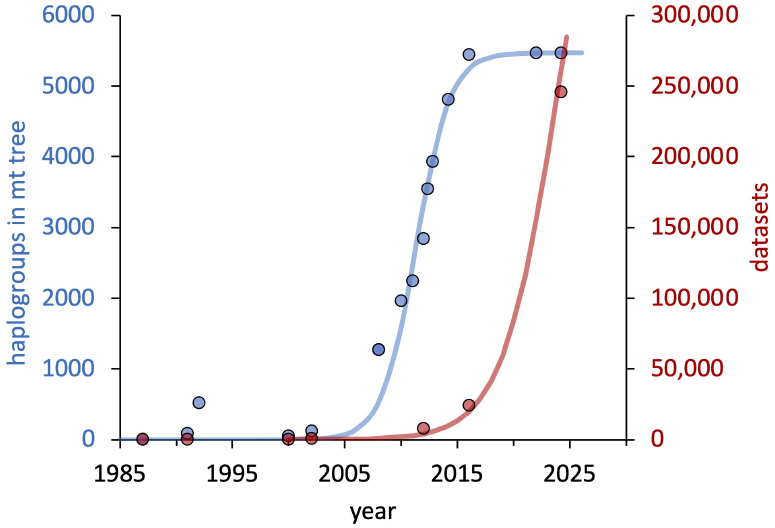
The growth of the mtDNA haplotree (blue) and number of consumer full-chromosome tests (red).

In 2020, FTDNA launched its Million Mito Project with the promise that “The more people who take the mtFull Sequence test (mtDNA), the more branches, twigs, and leaves can be discovered and mapped.” It’s a laudable goal and has had enthusiastic support from blogger Roberta Estes:. But are its expectations realistic?

The fundamental reason for the stalling of the mt tree lies in the numbers adduced here: with an effective length of only 4800 basepairs (3000-fold smaller than the measured Y chromosome), usable mutations are rare events that occur on average only every 2900 years (30-fold less often than Y SNPs). This means that new mtDNA testers will tend to pile up in haplogroups with common ancestry about 2900 years ago because there just aren’t more recent mutations to pick the groups apart. This is exactly what we see in a survey of all the leaf nodes (= terminal haplogroups) of the two haplotrees, shown in Figure 21, where the upper table gives average values of tMRCA, number of DNA testers, and SNPs for all leaf nodes. The lower chart shows how very different Y and mtDNA are in their allocation of information-per-tester.

**Figure 21.**
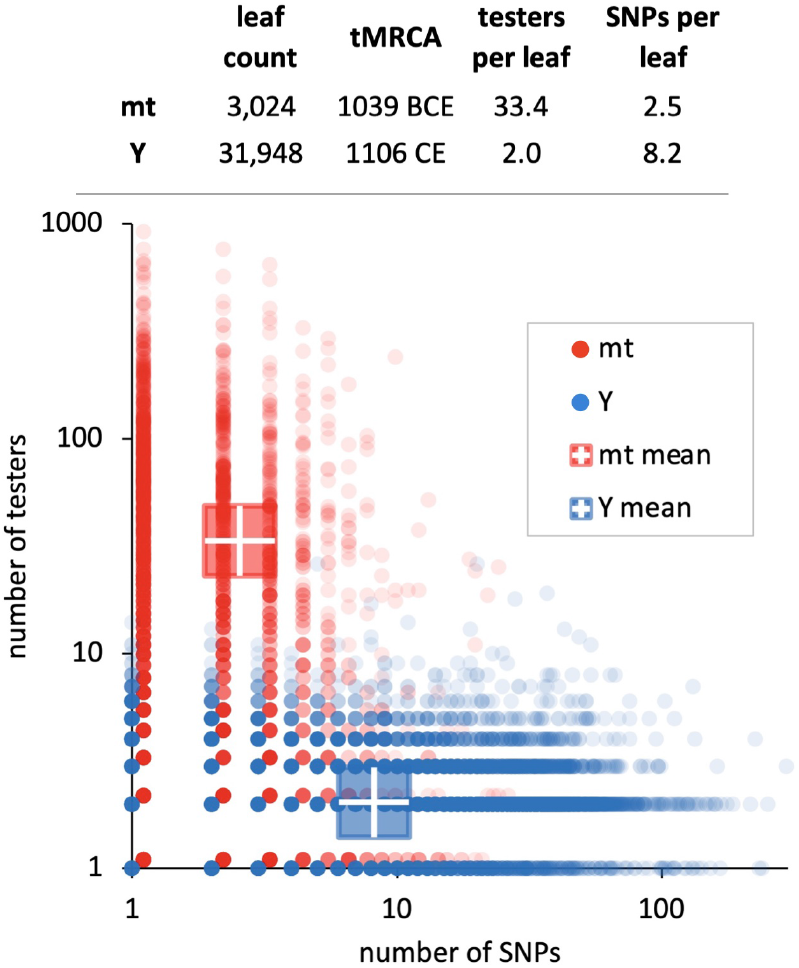
Upper table: Statistics for leaf nodes of the mt and Y haplotrees. Figure: Each dot is a leaf node (terminal haplogroup) in the mt(red) or Y(blue) haplotree. The blue cross shows, for example, that in Y DNA a leaf node averages two testers and 8 mutations.

With an average of 33 testers and 2.5 SNPs per leaf node, leaves (on average) might in principle be split into 2-3 haplogroups with tMRCAs closer to the present. The same might be done throughout the tree for the 20% of nodes with 3 or more SNPs. The fact that this hasn’t yet happened suggests that it’s inherently rare (due the linkage produced by the editing model above) and/or difficult in the tree-imputation process. The lessons learned from Y short tandem repeat (STR) mutations (which, like mtDNA SNPs, are oversampled within a bounded range) will apply; specifically, saturation (13,000 mutations piled onto 4800 locations) means that many “back mutations” (reversions) and “parallel mutations” (repetitions) – have and will continue to occur. These make branch location uncertain, prone to error, and prone to produce “convergence” artifacts similar to those seen for STRs in which two DNA testers appear to be much more closely related than they really are.

What might be done? Haplotrees are commonly inferred by maximum parsimony or maximum likelihood algorithms^19^. Some authors of published mt haplotrees simply ignore the most-repeated mutations, either for simple convenience or in the mistaken belief that these are anomalous “hot spots” instead of members of a continuous and self-consistent distribution: How many of the left-most mutations of Figure 16 should be ignored? Better perhaps might be a weighting scheme that utilizes all mutation data but gives more weight to those less frequently seen, but this would require experimental parameterization and may not be available in established software packages. It might also be possible to infer more mutations that are current invisible because they revert an earlier marker, though this poses a delicate balance between observed data and seductive over-parameterization.

The criteria for splitting a haplogroup could also be relaxed. Dür *et al* 2021^20^ do just this – a haplogroup in Phylotree build 17 (the 2016 archetype) is split if two or more testers have three or more mutations in common, none of which is present in the parent haplogroup. This adds a welcome 966 additional nodes (18%) to the tree, across all major haplogroups. However the authors do not mention reversions or repeats, which is worrisome since two testers might have a mutation in common because it is very frequent, instead of being inherited. In any case this tree is seldom cited and its database has been stagnant since 2019, so perhaps this stands as a beta-version, a predictor rather than a released and validated output.

Certainly jackknifing or bootstrapping (to create alternative trees with subsets of the data) will be in order to establish some sort of confidence level for branch assignments.

Efforts to make the mtDNA haplotree wider (more branching throughout) and deeper (more leaf nodes with more recent tMRCAs) would be very welcome, but it hasn’t happened in nine years despite an order of magnitude more datasets. Recomputing the tree with private variants is perhaps the most straightforward approach; relaxation of haplogroup splitting criteria could also be considered, though with attention to the misassignments likely with high numbers of repetitive mutations.

## Methods

### Assignment of tMRCAs to haplogroups

tMRCAs are assigned to all mtDNA haplogroups in two stages: interpolation from the less complete set published values, followed by linear scale adjustment to fit the distribution of ancient DNA evidence. Interpolation is from table S5 of Behar *et al* 2012^2^. tMRCAs for unassigned SNPs are pro-rated by mutation counts between SNPs with assigned dates; winsorized means of SNP counts are used for branched paths. Interpolation on branched paths uses data from all branches; details in source code below.

### Calibration from Ancient DNA

The calibrating distributions the FAR statistic^7^ of Figure 5 require ancient DNA samples with both radiocarbon dates and haplogroup assignments, which are sourced from the collection of [Carlos Quiles](https://indo-european.eu/ancient-dna/) and the Reich lab [Allen Ancient DNA Resource](https://dataverse.harvard.edu/dataset.xhtml?persistentId=doi:10.7910/DVN/FFIDCW). The “sliding” of the mt haplotree in Figure 4 is done by binary search, varying the root tMRCA until the mt distribution shows the same fraction with FAR *<* 1 as Y DNA from the same ancient samples.

### Years Between Mutations

Counting years between haplogroups (Figures 9, 10) is done by recursive haplotree traversal; the result relies on the complete assignment of tMRCAs in this work. While averaging may be done across the haplotree, the distribution of inter-mutation time is measurable only for unit segments, namely parent-child pairs without any intervening unplaced mutations.

### Counting Replicate Mutations

The results of Figures 14, 15 are achieved by haplotree traversal (source code below). This has apparently not been done before for mtDNA, or previous authors would have noted that particular loci are not unusual “hot spots” but rather part of a smooth distribution.

### Power Law Distributions

Estimation of the exponent *α* of Figures 16, 17 is done by Nelder-Mead fitting^21^ to the rank-frequency data, with error estimates from the curvature at the minimum.

### Current Haplotrees

FTDNA maintains the world’s most complete haplotrees which are accessible as large JSON files for [mtDNA](https://www.familytreedna.com/public/mt-dna-haplotree/get) and [Y DNA](https://www.familytreedna.com/public/y-dna-haplotree/get) and also as [smaller minimal files](https://scaledinnovation.com/gg/treeExplorer.html?getTrees).

### Source code

Source code is available for [specific routines](https://scaledinnovation.com/gg/reports/mt2/mtMethods.html) as well as [general](https://scaledinnovation.com/gg/gg.js) and [mathematical](https://scaledinnovation.com/gg/mathLib.js) utilities. Source code for the [mtDNA mutation simulation](https://scaledinnovation.com/gg/mtDNAbagrowModel.html) is available in that stand-alone application. Other calculations (histograms, distributions) are done in Excel.

## References

1. General references for mitochondrial population genetics Phylotree’s extensive bibliography Cann, R.L., Stoneking, M. and Wilson, A.C., 1987. Mitochondrial DNA and human evolution. Nature, 325(6099), pp.31–36. Sykes, B., 2010. The seven daughters of Eve: The science that reveals our genetic ancestry. WW Norton & Company. Hedges, S.B., Kumar, S., Tamura, K. and Stoneking, M., 1992. Human origins and analysis of mitochondrial DNA sequences. Science, 255(5045), pp.737–739. Ingman, M., Kaessmann, H., Pääbo, S. and Gyllensten, U., 2000. Mitochondrial genome variation and the origin of modern humans. Nature, 408(6813), pp.708–713. Di Rienzo, A. and Wilson, A.C., 1991. Branching pattern in the evolutionary tree for human mitochondrial DNA. Proceedings of the National Academy of Sciences, 88(5), pp.1597–1601. Templeton, A., 2002. Out of Africa again and again. Nature, 416(6876), pp.45–51. Kivisild, T. Maternal ancestry and population history from whole mitochondrial genomes. Investig Genet 6, 3 (2015). Hernández, C.L., 2023. Mitochondrial DNA in Human Diversity and Health: From the Golden Age to the Omics Era. Genes, 14(8), p.1534. Översti, S. and Palo, J.U., 2022. Variation in the substitution rates among the human mitochondrial haplogroup U sublineages. Genome Biology and Evolution, 14(7), p.evac097. Van Oven, M. and Kayser, M., 2009. Updated comprehensive phylogenetic tree of global human mitochondrial DNA variation. Human mutation, 30(2), pp.E386–E394. Zheng, HX., Yan, S., Qin, ZD. et al. MtDNA analysis of global populations support that major population expansions began before Neolithic Time. Sci Rep 2, 745 (2012). Connell, J.R., Benton, M.C., Lea, R.A. et al. Pedigree derived mutation rate across the entire mitochondrial genome of the Norfolk Island population. Sci Rep 12, 6827 (2022). Connell finds µ = 5.8×10^−9^ including heteroplasmy (difference within the same individual) and 1.3×10^−8^ excluding heteroplasmy. Maier, P.A., Runfeldt, G., Estes, R.J. et al. 2022. African mitochondrial haplogroup L7: a 100,000-year-old maternal human lineage discovered through reassessment and new sequencing. Sci Rep 12, 10747.

2. Behar, D.M., Van Oven, M., Rosset, S., Metspalu, M., Loogväli, E.L., Silva, N.M., Kivisild, T., Torroni, A. and Villems, R., 2012. A “Copernican” reassessment of the human mitochondrial DNA tree from its root. The American Journal of Human Genetics, 90(4), pp.675–684.,

3. Matching haplogroups between data sources that use the lineage-based system of Richards^6^ (i.e. U3a2a1a) carries some risk since established names may change if new branches are found; this problem does not exist for the mutation-based nomenclature that is increasingly common in the Y haplotree. In this case it is fortunate that the mt tree has been growing very slowly.

4. Adamov, D. et al. 2015, Defining a New Rate Constant for Y-Chromosome SNPs based on Full Sequencing Data, Russian Journal of Genetic Genealogy 7:68–89 The mutation count for any given lineage may be averaged either by leaf node or by branches. The former treats all DNA testers equally but is strongly influenced by European ancestry testing bias. Averaging by branch is less biased but more prone to statistical variance when a sub-branch has very few testers. The whole-tree difference is 3-7%: for Y DNA the number of mutations from the root A-PR2921 to the present day is 1955 by leaf-averaging and 2584 by branch-averaging. For mtDNA from RSRS, the values are 55.8 for leaf-averaging and 49.4 for branch averaging. Behar (2012) gives an mtDNA average to RSRS of 57.1. We can use 54 as a consensus value.

5. Mishmar, D., Ruiz-Pesini, E., Golik, P., Macaulay, V., Clark, A.G., Hosseini, S., Brandon, M., Easley, K., Chen, E., Brown, M.D., et al. (2003). Natural selection shaped regional mtDNA variation in humans. Proc. Natl. Acad. Sci. USA 100, 171–176. Soares, P., Ermini, L., Thomson, N., Mormina, M., Rito, T., Röhl, A., Salas, A., Oppenheimer, S., Macaulay, V. and Richards, M.B., 2009. Correcting for purifying selection: an improved human mitochondrial molecular clock. The American Journal of Human Genetics, 84(6), pp.740–759. Cabrera, V.M., 2021. Human molecular evolutionary rate, time dependency and transient polymorphism effects viewed through ancient and modern mitochondrial DNA genomes. Scientific Reports, 11(1), p.5036. Rieux, A., Eriksson, A., Li, M., Sobkowiak, B., Weinert, L.A., Warmuth, V., Ruiz-Linares, A., Manica, A. and Balloux, F., 2014. Improved calibration of the human mitochondrial clock using ancient genomes. Molecular biology and evolution, 31(10), pp.2780–2792. Fu, Q., Mittnik, A., Johnson, P.L., Bos, K., Lari, M., Bollongino, R., Sun, C., Giemsch, L., Schmitz, R., Burger, J. and Ronchitelli, A.M., 2013. A revised timescale for human evolution based on ancient mitochondrial genomes. Current biology, 23(7), pp.553–559.

6. Richards, M.B., Macaulay, V.A., Bandelt, H.J. and Sykes, B.C., 1998. Phylogeography of mitochondrial DNA in western Europe. Annals of human genetics, 62(3), pp.241–260.

7. FAR = formed-to-age ratio. The formed date is when the SNP first appeared, and the age date is the radiocarbon date of the DNA sample, with both dates as years-before-present. An ancient sample has FAR ≫1 if the person lived very soon after the SNP first appeared, and FAR ≈1 if they lived considerably later. Note that SNP formation dates are not directly measurable but may be inferred from tMRCAs. At some point after a SNP appears, a person with that SNP will have two or more children each with modern descendants who have done DNA testing. From those tests we can infer the time to that branch-point, which is the SNP’s tMRCA. In a rapidly expanding population with many surviving lineages, tMRCA and formation are very close. But for older and leaner lineages, a SNP may appear long before one of the originator’s descendants has two surviving lineages, and additional separate mutations may occur in that time. The convention at YFull is to assign a SNP’s formation date to its parent SNP’s tMRCA. But the expected error is least if the formation date is halfway between the tMRCAs of a SNP and its parent, which is my practice.

8. We have to be careful with terminology. Markers on the Y haplotree are named by mutation, for example in E-SK1027 the primary haplogroup is E and SK1027 denotes a particular mutation, so it’s easy to refer (incorrectly) to E-SK1027 as a SNP (a mutation) instead of as a haplogroup. 93% of Y mutations have been seen only once, so referring to E-SK1027 as a SNP does not cause much confusion. But mtDNA uses lineage nomenclature, so for example haplogroup U6c2 is named by its inferred position in the tree, under U6c and sibling to U6c1. U6c2 is placed there by the mutation C194T – but C194T is not synonymous with U6c2 because it also appears in 13 other contexts.

9. McDonald, I., 2021. Improved models of coalescence ages of Y-DNA haplogroups. Genes, 12(6), p.862.

10. Here SNPs = mutations = haplogroups because we’re only counting haplogroups on the Y tree that differ by one SNP. There are many inter-haplogroup branches that contain multiple SNPs, but if we included those we’d be implicitly averaging the very number that we want to measure. So Figures 9 and 10 are both limited to single-mutation segments of the Y and mt haplotrees, in order to see the un-smoothed distribution of these mutation arrival times. The initial droop in the Y distribution may reflect ad-hoc adjustments by FTDNA in their tMRCA assignments; to avoid child-older-than-parent cases they may apply rules to haplogroups that are close in time. Regardless, given the complexity of tMRCA estimation the fit to an exponential is excellent.

11. Half of these mutations occur in the third position of the codon, the wobble base, as expected. To a first approximation such synonymous mutations are silent and not under selection pressure. However some may nonetheless be disfavored, for example if the matching transfer RNA or aminoacyl-tRNA synthetase were scarce, then translation of the mutant codon could stall on the ribosome.

12. Each mitochondrion may have 1 to 5 copies of the genome and a typical vertebrate cell may have 100 to 10,000 mitochondria. Mature oocytes may have 100,000 mitochondria and 150,000 copies of the mt genome Filograna, R., Mennuni, M., Alsina, D. and Larsson, N.G., 2021. Mitochondrial DNA copy number in human disease: the more the better?. FEBS letters, 595(8), pp.976–1002. T. Wai, A. Ao, X. Zhang, D. Cyr, D. Dufort, E.A. Shoubridge The role of mitochondrial DNA copy number in mammalian fertility Biol. Reprod., 83 (2010), pp.52–62, 10.1095/biolreprod.109.080887 Mitophagy is a specific process for culling damaged mitochondria (Wang, S., Long, H., Hou, L., Feng, B., Ma, Z., Wu, Y., Zeng, Y., Cai, J., Zhang, D.W. and Zhao, G., 2023. The mitophagy pathway and its implications in human diseases. Signal Transduction and Targeted Therapy, 8(1), p.304.) and it is not difficult to suppose that even a small drop in efficiency (due to a mutation) would remove that mtDNA from the pool. Given the high intracellular numbers and independence of mitochondrial reproduction, that process could take place continuously, independently, and much faster than selection of the host organism’s nuclear DNA. mtDNA goes through at least two bottlenecks (dramatic reductions in copy number) in oogenesis and early embryogenesis, which could be points for strong purifying selection: Mishra, P. and Chan, D.C., 2014. Mitochondrial dynamics and inheritance during cell division, development and disease. Nature reviews Molecular cell biology, 15(10), pp.634–646.

13. Deeparnab Chakrabarty, D. April 2023, Lecture notes, Department of Computer Science, Dartmouth College

14. In the simulations these values of alpha mean that for control DNA, 40% of the time a new mutation is allotted to any position at random and 60% to an already-mutated position in proportion to how many mutations are already there (the rich-get-richer generator). For coding DNA it’s 60%-40% the other way.

15. Newman, M.E., 2005. Power laws, Pareto distributions and Zipf’s law. Contemporary physics, 46(5), pp.323-351. and A. Clauset, C.R. Shalizi, and M.E.J. Newman, Power-law distributions in empirical data, SIAM Review 51(4), 661–703 (2009) Bagrow, J.P., Sun, J. and ben-Avraham, D., 2008. Phase transition in the rich-get-richer mechanism due to finite-size effects. Journal of Physics A: Mathematical and Theoretical, 41(18), p.185001., Newman, M.E.J. and Sneppen, K., 1996. Avalanches, scaling, and coherent noise. Physical Review E, 54(6), p.6226. Sneppen, K. and Newman, M.E., 1997. Coherent noise, scale invariance and intermittency in large systems. Physica D: Nonlinear Phenomena, 110(3-4), pp.209–222. Newman, M.E.J., 1996. Self-organized criticality, evolution and the fossil extinction record. Proceedings of the Royal Society of London. Series B: Biological Sciences, 263(1376), 1605-1610. Newman, M.E.J. and Palmer, R.G., 1999. Models of extinction: A review. Miller, G.A. 1957, Some effects of intermittent silence. American Journal of Psychology 70, 311–314.

16. Shokolenko, I. and Alexeyev, M., 2022. Mitochondrial DNA: consensuses and controversies. DNA, 2(2), pp.131-148. Falkenberg, F. 2018. Essays in Biochemistry, 62 287–296

17. Arnason, U., Xu, X. and Gullberg, A., 1996. Comparison between the complete mitochondrial DNA sequences of Homo and the common chimpanzee based on nonchimeric sequences. Journal of Molecular Evolution, 42, pp.145–152. Andreeva, T.V., Manakhov, A.D., Gusev, F.E., Patrikeev, A.D., Golovanova, L.V., Doronichev, V.B., Shirobokov, I.G. and Rogaev, E.I., 2022. Genomic analysis of a novel Neanderthal from Mezmaiskaya Cave provides insights into the genetic relationships of Middle Paleolithic populations. Scientific Reports, 12(1), p.13016. Reich, D., Green, R.E., Kircher, M., Krause, J., Patterson, N., Durand, E.Y., Viola, B., Briggs, A.W., Stenzel, U., Johnson, P.L. and Maricic, T., 2010. Genetic history of an archaic hominin group from Denisova Cave in Siberia. Nature, 468(7327), pp.1053–1060. Pozzi, L., Hodgson, J.A., Burrell, A.S., Sterner, K.N., Raaum, R.L. and Disotell, T.R., 2014. Primate phylogenetic relationships and divergence dates inferred from complete mitochondrial genomes. Molecular phylogenetics and evolution, 75, pp.165–183.

18. I have generally glossed over the distinction between counting mutations and counting mutatable positions. About 12% of mt mutations occur at the same position as another but substitute a different nucleotide; in the current FTDNA mt haplotree there are 4128 mutatable positions and 4626 observed mutations. Mutations are the relevant entities for genealogy, since they are distinct heritable markers despite 12% being at the same position in the chromosome.

19. Wikipedia Phylip Guindon, S. and Gascuel, O., 2003. A simple, fast, and accurate algorithm to estimate large phylogenies by maximum likelihood. Systematic biology, 52(5), pp.696–704.

20. Dür, A., Huber, N. and Parson, W., 2021. Fine-tuning phylogenetic alignment and haplogrouping of mtDNA sequences. International Journal of Molecular Sciences, 22(11), p.5747.

21. Press, W. H. 1989. Numerical recipes in Pascal: the art of scientific computing (Vol. 1). Cambridge university press. Nelder-Mead fitting is function amoeba.

